# Dynamic and stationary brain connectivity during movie watching as revealed by functional MRI

**DOI:** 10.1101/2021.09.14.460293

**Authors:** Xin Di, Zhiguo Zhang, Ting Xu, Bharat B. Biswal

**Author notes:** Corresponding author: Xin Di, Ph.D. 604 Fenster Hall, University Height, Newark, NJ, 07102, USA, Bharat B. Biswal, Ph.D. 607 Fenster Hall, University Height, Newark, NJ, 07102, USA.

## Abstract

Spatially remote brain regions show synchronized activity as typically revealed by correlated functional MRI (fMRI) signals. An emerging line of research has focused on the temporal fluctuations of connectivity; however, its relationships with stationary connectivity have not been clearly illustrated. We examined dynamic and stationary connectivity when the participants watched four different movie clips. We calculated point-by-point multiplication between two regional time series to estimate the time-resolved dynamic connectivity, and estimated the inter-individual consistency of the dynamic connectivity time series. Widespread consistent dynamic connectivity was observed for each movie clip, which also showed differences between the clips. For example, a cartoon movie clip, Wall-E, showed more consistent of dynamic connectivity with the posterior cingulate cortex and supramarginal gyrus, while a court drama clip, A Few Good Men, showed more consistent of dynamic connectivity with the auditory cortex and temporoparietal junction, which might suggest the involvement of specific brain processing for different movie contents. In contrast, the stationary connectivity as measured by the correlations between regional time series was highly similar among the movie clips, and showed fewer statistically significant differences. The patterns of consistent dynamic connectivity could be used to classify different movie clips with higher accuracy than the stationary connectivity and regional activity. These results support the functional significance of dynamic connectivity in reflecting functional brain changes, which could provide more functionally relevant information than stationary connectivity.

## 1. Introduction

The human brain exhibits a highly synchronized structure of activity as revealed by functional MRI (fMRI) in resting-state (Biswal et al., 1995, 2010), during task performance (Cole et al., 2014; Di et al., 2020; Krienen et al., 2014), and during watching naturalistic stimuli such as movies (O’Connor et al., 2017; Vanderwal et al., 2019). Functional connectivity, as measured by the correlations of observed blood-oxygen-level-dependent (BOLD) signals (Biswal et al., 1995; Friston, 1994), have been widely used to examine the organization of large-scale brain networks (Margulies et al., 2016; Salvador et al., 2005; Yeo et al., 2011) and to parcellate small brain structures such as the thalamus and striatum (Di Martino et al., 2008; Tian et al., 2020; Yuan et al., 2016). However, the spatial distribution of functional connectivity is highly similar across different tasks and movie watching conditions (Cole et al., 2014; Di et al., 2020; Krienen et al., 2014; Vanderwal et al., 2019). To localize functionally meaningful connections, it is therefore critical to examine the time-varying fluctuations of connectivity (Allen et al., 2014; Di and Biswal, 2020; Hutchison et al., 2013), as well as the changes of functional connectivity between different task conditions (Di and Biswal, 2019; Fornito et al., 2012; Friston et al., 1997).

Time-varying dynamic connectivity is mostly studied in the resting-state by using the sliding-window approach (Allen et al., 2014; Hutchison et al., 2013; Lurie et al., 2020). It has been shown that the variability of dynamic connectivity fluctuations is lower between regions from the same functional networks and higher between regions from different networks (Fu et al., 2017), resulting in an overall negative correlation with the stationary functional connectivity (Thompson and Fransson, 2015; Zhang et al., 2018). However, because of the unconstrained nature of the resting-state, it is difficult to ensure that the obtained dynamic connectivity estimates are functionally meaningful or simply resulting from noise (Lindquist et al., 2014). Until recently, dynamic connectivity is also studied when the participants were given complex stimuli, such as watching movie clips (Di and Biswal, 2020). The advantage of using a movie stimulus is that the time course of dynamic connectivity can be compared across participants. If there is high inter-individual similarity (Hasson et al., 2004; Nastase et al., 2019), then it may imply that the observed dynamic connectivity is functionally meaningful and is relevant to the processing of the stimuli.

In our previous study, we have demonstrated the inter-individual consistency of dynamic connectivity when different participants watched the same animated movie Partly Cloudy (Di and Biswal, 2020). By using a seed-based analysis, we identified highly consistent dynamic connectivity between the supramarginal gyrus and posterior cingulate gyrus, two regions that are critical in the processes of empathy and theory of mind (Richardson et al., 2018). Moreover, among a set of regions of interest, the dynamic connectivity pattern was largely dissociated with the stationary functional connectivity that was measured by the correlations of the time series from the entire run. For example, the stationary connectivity between the supramarginal gyrus and posterior cingulate gyrus was close to zero, while the windowed dynamic connectivity showed highly consistent fluctuations. To date, only a handful of studies have examined dynamic connectivity during movie watching (Cooper et al., 2021; Di and Biswal, 2020; Freitas et al., 2020; Simony and Chang, 2020). It is still largely unknown how the spatial pattern is modulated by different movie contents, and how dynamic connectivity is spatially distributed.

The commonly used stationary connectivity has also been employed on movie watching data. Two studies have investigated whole-brain stationary connectivity pattens during movie conditions, but they both found that the connectivity matrices were highly similar across different movies clips (Tian et al., 2021; Vanderwal et al., 2019). This suggests that the timescale where stationary connectivity is calculated may not be sensitive to the fast brain interactions and the changes of movie contents. On the other hand, dynamic connectivity may be more suitable to capture brain interactions in finer timescale, therefore is able to reflect different movie contents. Direct comparisons between the dynamic and stationary connectivity are therefore necessary.

The central goal of this study is to compare dynamic and stationary connectivity in the context of movie watching. In addition to the previously analyzed Partly Cloudy dataset (Richardson et al., 2018), we also analyzed the Healthy Brain Network Serial Scanning Initiative (HBN-SSI) dataset (O’Connor et al., 2017), where the same participants watched three different movie clips. The video clips were derived from different types of movies, ranging from a science fiction cartoon comedy, a science fiction action film, to a court drama. It is reasonable to expect that different brain systems are involved in the process of the different movie clips.

We systematically examine the relationships between dynamic and stationary connectivity in terms of their spatial distributions and context modulations. The economic theory of brain network organization has suggested that the maintenance of long-range between-system communications is costly, and long-range and between-system connectivity may be more dynamic and depend on task demands (Bullmore and Sporns, 2012). In line with this account, the dynamic connectivity between different functional systems are more variable than those within functional systems (Fu et al., 2017; Thompson and Fransson, 2015), and task-modulated connectivity is also likely to be observed between regions from different functional networks (Di and Biswal, 2019). Similarly, for the movie-watching data, we speculate that dynamic connectivity might take place between regions from different functional modules. In contrast, the stationary connectivity might tightly reflect the organizations of brain networks, i.e., higher stationary connectivity between regions from the same functional networks, and lower stationary connectivity between regions from different networks. The dissociation might result in different spatial patterns between the dynamic and stationary connectivity.

## 2. Materials and Methods

### 2.1. FMRI dataset

We analyzed two publicly available fMRI datasets when participants watched different movie clips, the Partly Cloudy dataset (Richardson et al., 2018) and the HBN-SSI dataset (O’Connor et al., 2017). For the Partly Cloudy dataset, we analyzed the adults’ data where they watched the animated movie “Partly Cloudy”. And for the HBN-SSI dataset, we analyzed the data when the same participants watched three different movie clips four times from different types of movies.

#### 2.1.1. Partly Cloudy dataset

The Partly Cloudy data were obtained through openneuro (https://openneuro.org/; accession #: ds000228). Consistent with our previous study, we only included the adult participants (original n = 33) (Di and Biswal, 2020). After dropping data due to large head motion (see below) and poor brain coverage, the effective sample included 29 participants (17 females). The mean and standard deviation of age were 24.6 years and 5.3, respectively (age range: 18 to 39 years). The original study was approved by the Committee on the Use of Humans as Experimental Subjects (COUHES) at the Massachusetts Institute of Technology.

During the fMRI scan, the participants watched a 5.6-minute long silent version of Partly Cloudy (Pixar, 2009). MRI images were acquired on a 3-Tesla Siemens Tim Trio scanner with the standard Siemens 32-channel head coil. Blood-oxygen-level dependent (BOLD) sensitive fMRI images were collected with a gradient-echo EPI sequence in 32 interleaved near-axial slices (EPI factor: 64; TR: 2 s, TE: 30 ms, and flip angle: 90°). The participants were recruited for different studies with slightly different voxel sizes and slice gaps. Three participants had 3.13 mm isotropic voxels with no gap, and 26 participants had 3.13 mm isotropic voxels with a 10% gap. All the functional images were resampled to 3 mm isotropic voxel size during preprocessing. 168 functional images were acquired, with four dummy scans before the real scans to allow for steady-state magnetization. T1-weighted structural images were collected in 176 interleaved sagittal slices with 1 mm isotropic voxels (GRAPPA parallel imaging, acceleration factor of 3; FOV: 256 mm). More information can be found in Richardson et a. (2018).

#### 2.1.2. HBN-SSI dataset

The HBN-SSI dataset was obtained through the project website (http://fcon_1000.proiects.nitrc.org/indi/hbn_ssi/). Thirteen participants were recruited in the study. After removing data of four participants due to excessive head motion in any of the movie-watching sessions, data from 9 participants (four females) were included in the current analysis. All the participants are right-handed. The age range was from 23 to 37 years old (*Mean* = 29.4; SD = 5.5).

We selected the movie watching scans of three movie clips, Wall-E (Walt Disney Productions, 2008), The Matrix (Warner Bros., 1999), and A Few Good Men (Columbia Pictures, 1992), from the 12 repeated scanning sessions. Each clip is 10 minutes long. The three clips are from different movie genres with complete dramatic events. The Wall-E clip is from the beginning of the movie, where it sets up the background and world of the movie. It contains actions of the singe robot actor, Wall-E. There are no conversations, but only music in the background. The Matrix clip includes the segments when Neo first met with Morpheus and then awaken in the real world. The A Few Good Men clip is near the end of the movie, which mainly includes conversations and arguments in the courtroom. Each movie clip was watched by the same participant four times in separate sessions, and the order of the clips was counterbalanced across sessions. The fMRI data were scanned using an EPI sequence with the following parameters, TR: 1,450 ms, TE: 40 ms, flip angle: 55°, and voxel size: 2.46 × 2.46 s 2.5 mm^3^ without any gap. Four hundred and twenty images were scanned for each run. However, for one participant, there were only 410 images for several sessions. We, therefore, used the first 410 images for the current analysis for all the subjects and sessions.

Lastly, the MPRAGE image from the first sequential scanning session of each participant was used to assist preprocessing of the fMRI data. The scanning parameters include TR, 2,730 ms; TE 1.64 ms; flip angle, 7°; voxel size 1 × 1 × 1 mm^3^ with no gap. More information about the study design and MRI acquisitions can be found in O’Connor et al., (2017).

### 2.2. FMRI data preprocessing

The fMRI data preprocessing was performed by using SPM12 (SPM, RRID: SCR_007037) under MATLAB environment (https://www.mathworks.com/). The two datasets were preprocessed using very similar pipelines. Specifically, the anatomical image of each participant was first segmented into gray matter, white matter, cerebrospinal fluid, and other tissue types, and normalized into standard Montreal Neurological Institute (MNI) space. The functional images of each session and subject were aligned to the first image of their specific session and were coregistered to the skull stripped anatomical image of the subject. The deformation field maps obtained from the segmentation step were used to normalize all the functional images into MNI space. The fMRI images from the Partly Cloudy dataset were resampled to 3 × 3 × 3 mm^3^ voxel size; and the images from the HBN-SSI dataset were resampled to 2.5 × 2.5 × 2.5 mm^3^, which were chosen according to their respective original voxel sizes. All the functional images were then spatially smoothed using an 8 mm Gaussian Kernel. Lastly, we defined a generalized linear model (GLM) for each session and subject by using 24 head motion variables (Friston et al., 1996) and a constant term as regressors, with implicit high-pass filtering at 1/128 Hz. After model estimation, the residual images were saved for further analysis.

We calculated framewise displacement for translation and rotation for each session and participant (Di and Biswal, 2015). For the Partly Cloudy dataset, we used strict criteria of maximum framewise displacement of 1.5 mm or 1.5° to discard data with large head movements. Two participants’ data were discarded accordingly. For the HBN-SSI dataset, a participant’s data were discarded if any of the sessions exceeded the criteria. We adopted a slightly liberal criterion of maximum framewise displacement greater than 2.5 mm or 2.5° or mean framewise displacement greater than 0.2 mm or 0.2°. Four participants’ data were removed accordingly.

### 2.3. Independent component analysis

Because the main goal of the current study is to study connectivity across brain regions, we adopted spatial independent component analysis (ICA) to define connectivity nodes. We extracted 20 and 80 ICs to represent different spatial scales of brain networks. We first analyzed the local activity and connectivity with 20-IC solutions to identify statistically significant local effects. We then calculated connectivity using the 80-IC solutions to examine their spatial distributions. For spatial ICA, the number of ICs that could be extracted depends on the number of time points for each participant/session. Theoretically, *t −1* components can be extracted where *t* represents the total number of time points. We chose 80, which is roughly half of the time points for the Partly Cloudy dataset.

Group ICA of fMRI Toolbox v3.0b (Group ICA of fMRI Toolbox, RRID: SCR_001953) was used for ICA (Calhoun et al., 2001). The ICA was performed for the Partly Cloudy and HBN-SSI datasets separately. After extraction, we manually selected the ICs that were related to functional networks and discarded the noise-like ICs. For the Partly Cloudy dataset, 16 and 65 ICs were considered functionally meaningful ICs for the 20-IC and 80-IC solutions, respectively. And for the HBN-SSI dataset, 16 and 54 ICs were kept. After ICA, the time series of each IC for each subject and session were back reconstructed by using the group ICA algorithm. The time series were used for further activity and connectivity analyses.

### 2.4. Inter-individual consistency of regional activity

For both regional activity and dynamic connectivity, we estimated the inter-individual consistency across participants and sessions. Conventionally, the intersubject correlation was used to study the inter-individual consistency (Chen et al., 2016; Hasson et al., 2004; Nastase et al., 2019). In our recent study, we have shown that the principal component analysis (PCA) can be used to estimate the inter-individual consistency, which is quantitively similar to intersubject correlation (Di and Biswal, 2021). Specifically, for a given region we have a *t* (# of time points) by *n* (# of participants) matrix *X. X* is a 168 × 29 matrix for the Partly Cloudy dataset and a 410 × 36 matrix (36 = 9 participants x 4 sessions) for each of the three movie clips from the HBN-SSI dataset. We performed PCA on the matrix *X* and obtained the percent variance explained by the first PC as a measure of intersubject consistency.

For the HBN-SSI dataset, there were four sessions for each participant and each movie. Ideally, the multi-session and multi-participant design can be used to differentiate the consistent and idiosyncratic responses. We have explored this issue and found that the within-participant consistency was mainly driven by the overall across-participant consistency, but not participant-specific idiosyncratic responses (see Supplementary materials section S1). Moreover, the idiosyncratic responses are not the focus of the current study. Therefore, we treated session and participant as separate data and calculated inter-individual consistency across all the sessions and participants.

### 2.5. Dynamic connectivity

The sliding-window approach is the most commonly used method to estimate dynamic connectivity (Allen et al., 2014; Di and Biswal, 2020; Fu et al., 2014). A recent development is to utilize point-by-point multiplications of two time series to approximate their dynamic connectivity, a.k.a. edge-centric time series (Faskowitz et al., 2020). The development comes from the intuition that the commonly used measure of functional connectivity, i.e., Pearson’s correlation coefficient, is the summation of the point-by-point multiplications of two z transformed variables divided by the sample size minus 1. Therefore, if we keep the original point-by-point multiplication time series, it can reflect estimates of dynamic connectivity at every time point. In Figure 1, we show the averaged time series of two networks, i.e., the posterior cingulate network and supramarginal network from the Partly Cloudy dataset. The averaged sliding-window and point-by-point multiplication time series were also shown. Strong negative multiplication values can be seen when the two original time series have strong anti-phase co-fluctuations. Indeed, the peaks in the posterior cingulate network and supramarginal network represent the theory-of-mind and pain empathy events, respectively, as indicated by the original paper (Richardson et al., 2018). The point-by-point multiplication indicates strong negative connectivity during these events. In contrast, the sliding-window correlation can only reflect a smoothed trend of such interactions. The point-by-point multiplication approach can avoid overly smoothing the time series data as done by the sliding-window approach, therefore providing better interpretability of the results. On the other hand, the multiplication term may be noisier and more prone to physiological noises and head motion artifacts. But this may be less problematic for movie watching data, where we can estimate the consistent effects cross individuals. We have performed statistical analyses on the Partly Cloudy dataset, and confirmed that the point-by-point multiplication approach had better statistical sensitivity (see Supplementary materials section S3). Therefore, we adopted the point-by-point multiplication approach to estimate dynamic connectivity.

**Figure 1.**
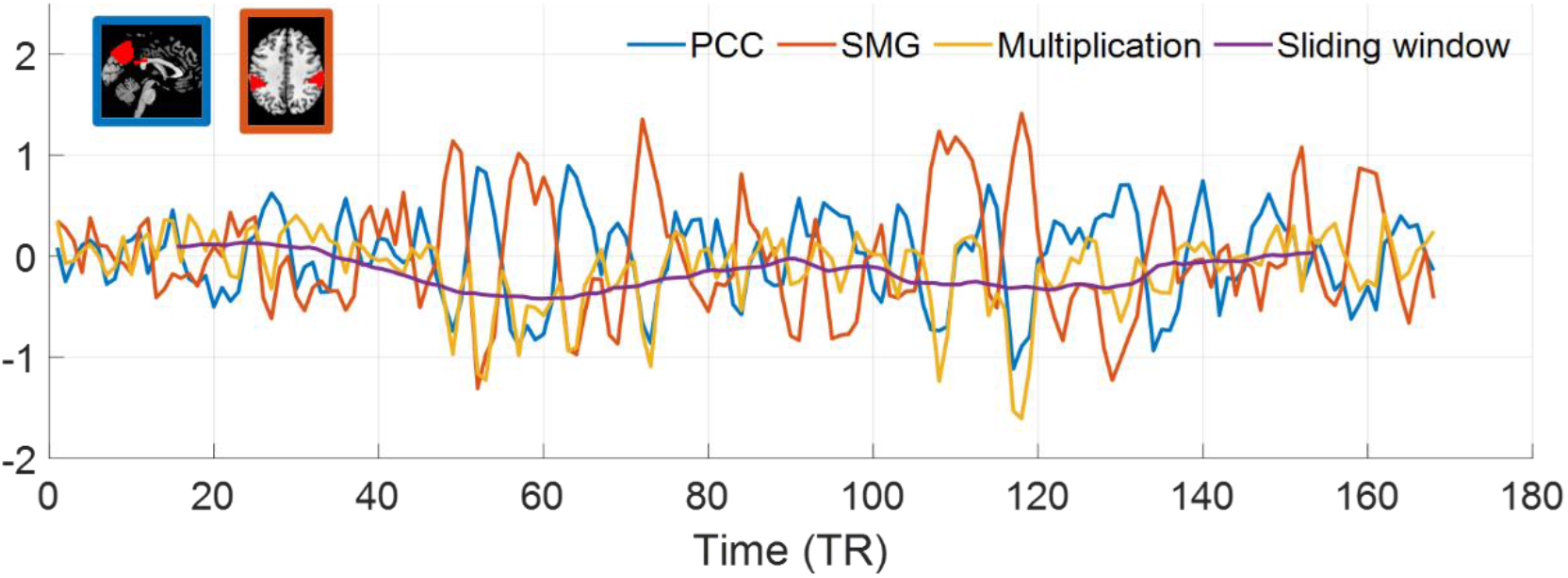
Averaged time series of regional activity in the posterior cingulate (PCC) network and supramarginal network (SMG), and their point-by-point multiplication and sliding-window dynamic connectivity when watching an animated short movie Partly Cloudy. The two brain networks shown in the inserted slices correspond to independent components #15 and # 6 in Figure 3, respectively.

We calculated the point-by-point multiplication between each pair of networks (ICs). The time series from each network (IC) were first z transformed, and then point-by-point multiplied. PCA was then performed on the multiplication time series across participants. To determine the statistical significance of the variance explained by the first PC, we performed circular time-shift randomization to determine the null distribution (Di and Biswal, 2021; Kauppi et al., 2010). The time series from the two network ICs from all the participants were circular-shifted with random delays. Point-by-point multiplications were then calculated for each participant, and PCA was performed. The randomization was performed 10,000 times for each pair of networks from the 16 networks. The real values were compared with the null distribution to perform statistical inferences. This resulted in a 16 × 16 matrix. False discovery rate (FDR) correction was used to correct for multiple comparisons (120: 16 × 15 / 2).

For the HBN-SSI dataset, we also compared the differences in variance explained by the first PC among the three movies. For a pair of networks ICs, the multiplication between two IC time series were first calculated, forming a 410 (time point) x 36 (participant/session) matrix for each movie. The matrices from the three movie clips were concatenated to a 410 × 108 matrix, and permutation was performed along the individual/session dimension to define three permutated matrices. The differences in variance explained by the first PC between each movie clip and the other two clips were calculated and compared with the permutated distributions of 10,000 times. FDR correction at p < 0.05 was used to correct multiple comparisons of all three movie clips.

The randomization-based statistics were performed for all the analyses in the 20-IC solutions. For the 80-IC solution, the goal of the analyses was not to identify specific statistically significant connections. Rather, we examined the spatial distributions of the dynamic connectivity, and their relations to stationary connectivity.

### 2.6. Relations to stationary connectivity

Next, we examined how the spatial distribution of consistent dynamic connectivity is associated with stationary connectivity. Here we focused on the finer spatial scale of 80-IC solutions. To assess the stationary connectivity, we calculated Fisher’s z transformed Pearson’s correlations across the included networks (ICs). The matrices were averaged across individuals, and transformed back to r quantities. First, we examined whether the dynamic and stationary connectivity has similar spatial distributions. For both matrices, the upper triangular part was converted to vectors, which were in turn correlated with each other between different movie clips. We adopted Spearman’s correlation coefficients to avoid violations of Gaussian distributions of the matrix data.

Next, we calculated connectivity gradients (Margulies et al., 2016; Vos de Wael et al., 2020) based on the stationary connectivity patterns in the HBN-SSI dataset. The gradients represent the extent to which nodes are similarly connected in the affinity matrix defined by the ICs. By calculating gradients, the brain networks (ICs) can be placed into a 2-D space based on their relative stationary connectivity strengths. The 2-D gradients reflect large-scale brain organizations between unimodal networks and higher-order transmodal areas (e.g. the default mode network) and between visual and sensorimotor regions (Margulies et al., 2016). We can next display dynamic connectivity in the 2-D space to illustrate whether the dynamic connectivity takes place between proximal or distal regions in the 2-D space. Specifically, we first calculated the gradients for each movie clip based on the group stationary connectivity matrices using the BrainSpace toolbox with diffusion mapping method (Vos de Wael et al., 2020). This was achieved in 3 steps. First, row-wise threshold at top 10% was performed on the stationary connectivity (i.e., leaving top 10% connections for each IC) to build the affinity matrix using the cosine similarity. After that, the normalized Laplacian matrix was calculated with alpha = 0.5 to build the transition matrix of a Markov chain. Finally, the eigenvalues and eigenvectors were extracted to compute the gradients from the probability map of the diffusion process at the time (*t* = 0) along the stationary connectivity graph. After gradients were calculated for each movie clip, they were then aligned across the movie clips with Procrustes alignment. The first two aligned gradients were then obtained to construct the 2-D gradient space. The network ICs were plotted as nodes in the 2D gradient space. Lastly, the dynamic connectivity between ICs from the three movie clips was mapped on the 2-D layout.

### 2.7. Movie clips classification

In addition to univariate analysis, we also explored whether the dynamic connectivity, stationary connectivity, and regional activity can reliably reflect an individual’s movie-watching condition (Finn et al., 2015). The analysis was performed based on the 54 network ICs from the 80-IC solution. For each participant of the HBN-SSI dataset, we calculated a connectivity or activity measure for each movie clip, and also the corresponding connectivity or activity measure for the remaining 8 participants. We compared three types of measures. First, we calculated the inter-individual consistency of point-by-point multiplications. The individual’s consistency was calculated across the four sessions of the same movie clips. The consistency of the remaining participants was calculated across 32 participants/sessions. The lower diagonal of the matrices was converted into a 1,431 (54 × 53 / 2) by 1 vector to perform the classification analysis. Second, we calculated mean stationary connectivity for the individual (averaged across 4 sessions) and the remaining participants (averaged across 32 sessions). Third, we calculated the inter-individual consistency of the regional activity. Similarly, individual measures were calculated across the four sessions, and the remaining participants’ measures were calculated for the 32 participants/sessions. Each measure was a 54 by 1 vector, which was used for the classification analysis.

We used a winner-take-all algorithm to perform the movie clip classifications. To classify the individual measure’s state (movie clips), the individual’s measures were correlated with the remaining participants’ measures from the three movie clips. The movie clips with the highest correlation were considered as the predicted class. The classification was performed for each of the 9 participants and 3 movie clips, from which we calculated confusion matrices among the three movie clips and the overall classification accuracy of all the three movie clips. Because three movie clips were used for the classifications, the chance level accuracy is 33.33%. To determine statistical significance, we adopted a permutation procedure to randomly shuffle the predicted movie label 10,000 times.

## 3. Results

### 3.1. Dynamic and stationary connectivity in 20-IC solution

We first examined the dynamic connectivity by calculating point-by-point multiplications between each pair of 16 networks (IC) on the Partly Cloudy dataset (Figure 3a). We found widespread inter-individual consistent effects at p < 0.05 of FDR correction. We also applied the sliding-window approach, which only yielded statistically significant effects on six pairs of networks (Supplementary materials section S3). This suggests that the point-by-point multiplication approach has better sensitivity to detect dynamic connectivity than the sliding-window approach. Although widespread, higher inter-individual consistency was found mostly involving one network of the higher visual networks (IC# 2, 3, and 4 in Figure 3c). And more interestingly, high inter-individual consistent dynamic connectivity was also found between the supramarginal and default mode network (IC # 6 and 15), which was similar to our previous analysis using a different analytic approach. In contrast, the stationary connectivity showed different spatial patterns than the dynamic connectivity (Figure 3b). The networks with known functional relations, e.g., all the visual related networks (IC # 1, 2, 3, and 4), had higher stationary connectivity, which showed square-like structures along the diagonal.

**Figure 2.**
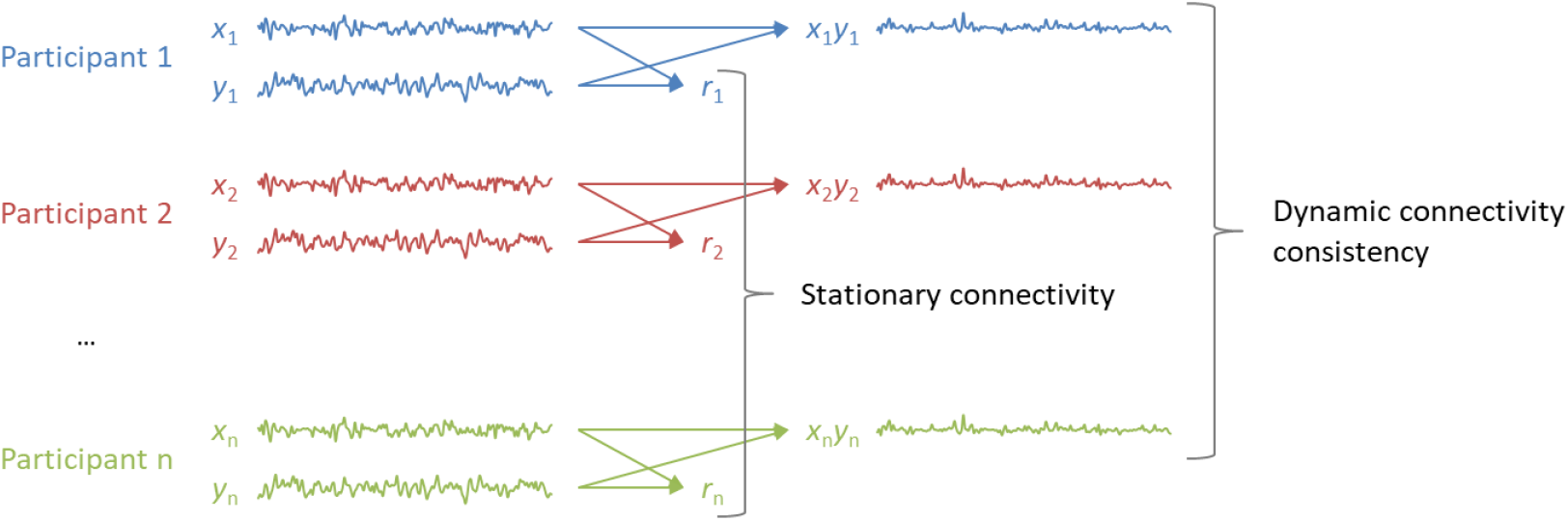
Illustration of the calculation of the consistency of dynamic connectivity and stationary connectivity.

**Figure 3.**
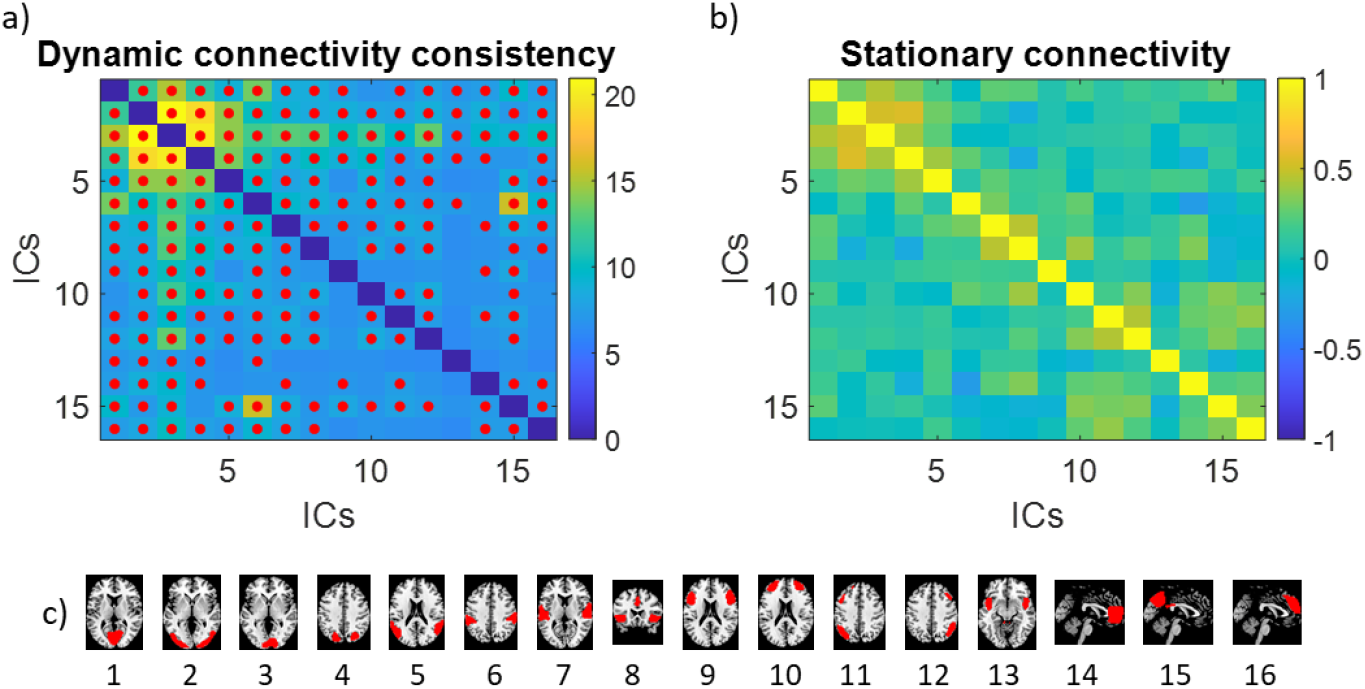
a) Inter-individual consistent point-by-point multiplications (dynamic connectivity) across the 16 networks (independent components, ICs) from the Partly Cloudy dataset. The colors in the matrix represent the percent variance explained by the first principal component of the point-by-point multiplication. The red dots indicate a false discovery rate (FDR) p < 0.05 using a circular time-shift randomization procedure. b) Mean stationary connectivity across the 16 network ICs. The networks are ordered roughly according to their functions. The locations of the networks are shown in c).

Figures 4a through 4c show the consistent point-by-point multiplications among the 16 networks (ICs) for the three movie clips in the HBN-SSI dataset. Consistent with the Partly Cloudy dataset, widespread consistent dynamic connectivity was observed. The networks that had more consistent dynamic connectivity were higher visual networks and the auditory network (IC 7). We further directly compared the consistency of dynamic connectivity among the three movies (Figure 4d through 4f). Compared with the other two movie clips, the Wall-E clip showed more consistent dynamic connectivity between the supramarginal network (IC 6) and many other networks, and between the posterior cingulate network (IC 13) and visual related networks. Compared with the other movie clips, the clip of The Matrix showed greater consistency between only three pairs of networks, among the posterior visual cortex, posterior parietal network, supramarginal network, and a left frontoparietal network. And lastly, compared with the other two movie clips, the A Few Good Men clip showed greater consistency in the multiplication of the auditory cortex (IC 7) with other networks, and between the temporoparietal junction network (IC 5) and other networks.

**Figure 4.**
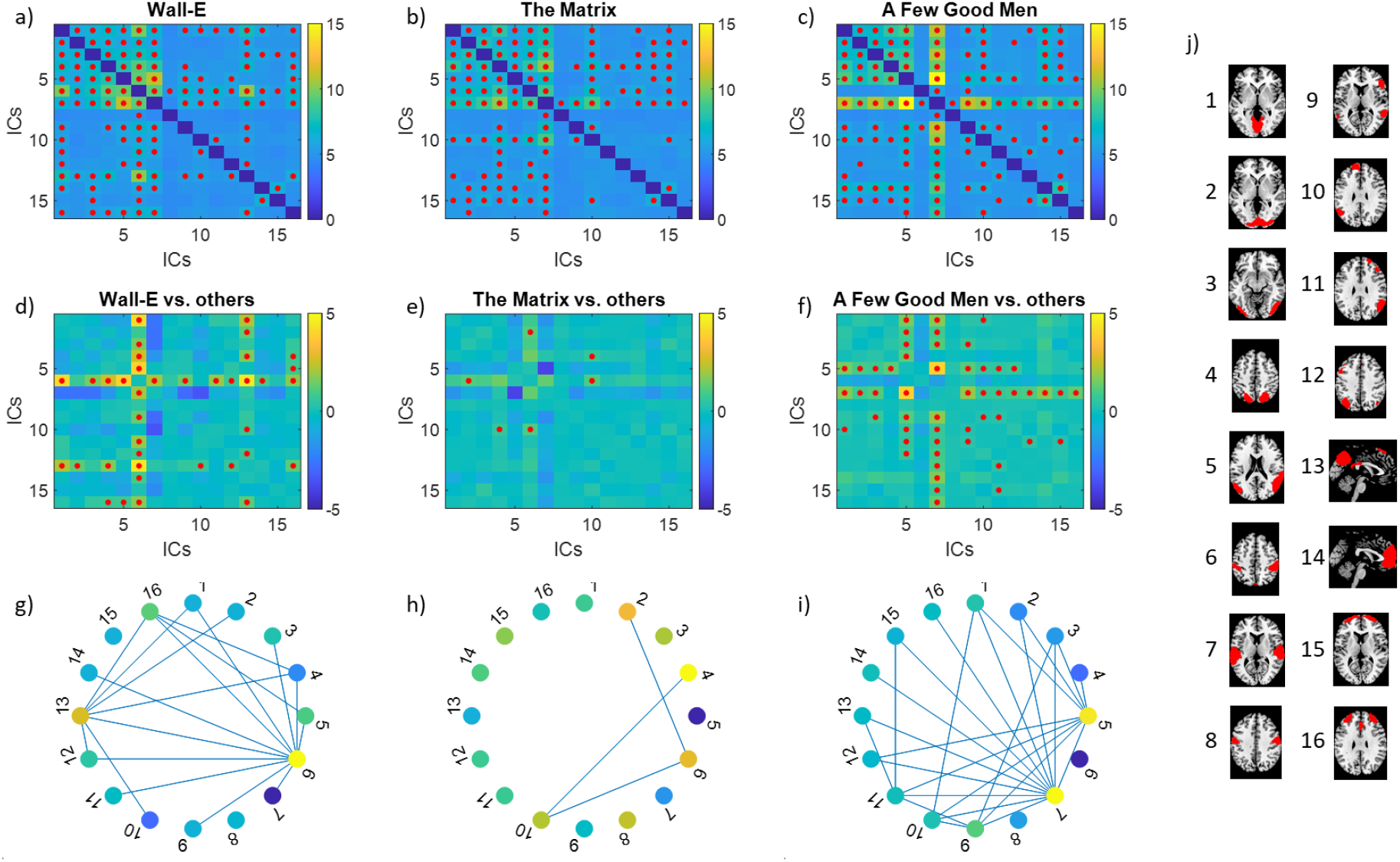
a) through c), consistency of dynamic connectivity across 16 networks (independent components, ICs) when watching the three movie clips in the Healthy Brain Network serial scanning initiative dataset. The colors in the matrices represent the percent variance explained by the first principal component of the point-by-point multiplication. d) through f), the differences in consistency of dynamic connectivity between each movie clip and the other two clips. The red dots indicate statistical significance at a false discovery rate p < 0.05 with permutation testing. The differences in consistency of dynamic connectivity are also shown in graph representations in g) through i), where the node color represents the consistency differences in regional activity. j) shows the representative maps for the 16 network ICs.

We also examined the relationships between regional activity consistency and dynamic connectivity consistency. We first compared the variance explained by the first PC for regional activity between the three movie clips (supplementary Figure S2). Four networks (ICs) showed higher inter-individual correlations in Wall-E compared with the other two movie clips, including the posterior cingulate cortex, supramarginal gyrus, left fronto-parietal, and medial and lateral prefrontal networks. Only one network covering the posterior parietal lobe showed higher inter-individual synchronization in The Matrix compared with the other two movie clips. Eight networks showed higher inter-individual correlations in A Few Good Men compared with the other movie clips, including the auditory cortex, medial visual, temporo-parietal junction, and a few fronto-parietal networks. More interestingly, the regions with greater consistency in regional activity in different movie clips seem to correspond well with the regions with many consistent dynamic connectivities (Figure 4g through 4i).

We next examined the stationary connectivity among the 16 networks (ICs) (Figure 5). Not surprisingly, the overall patterns in the three movies were very similar. When directly comparing the differences among the three movies, no statistically significant differences were found even at the p < 0.05 threshold.

**Figure 5.**
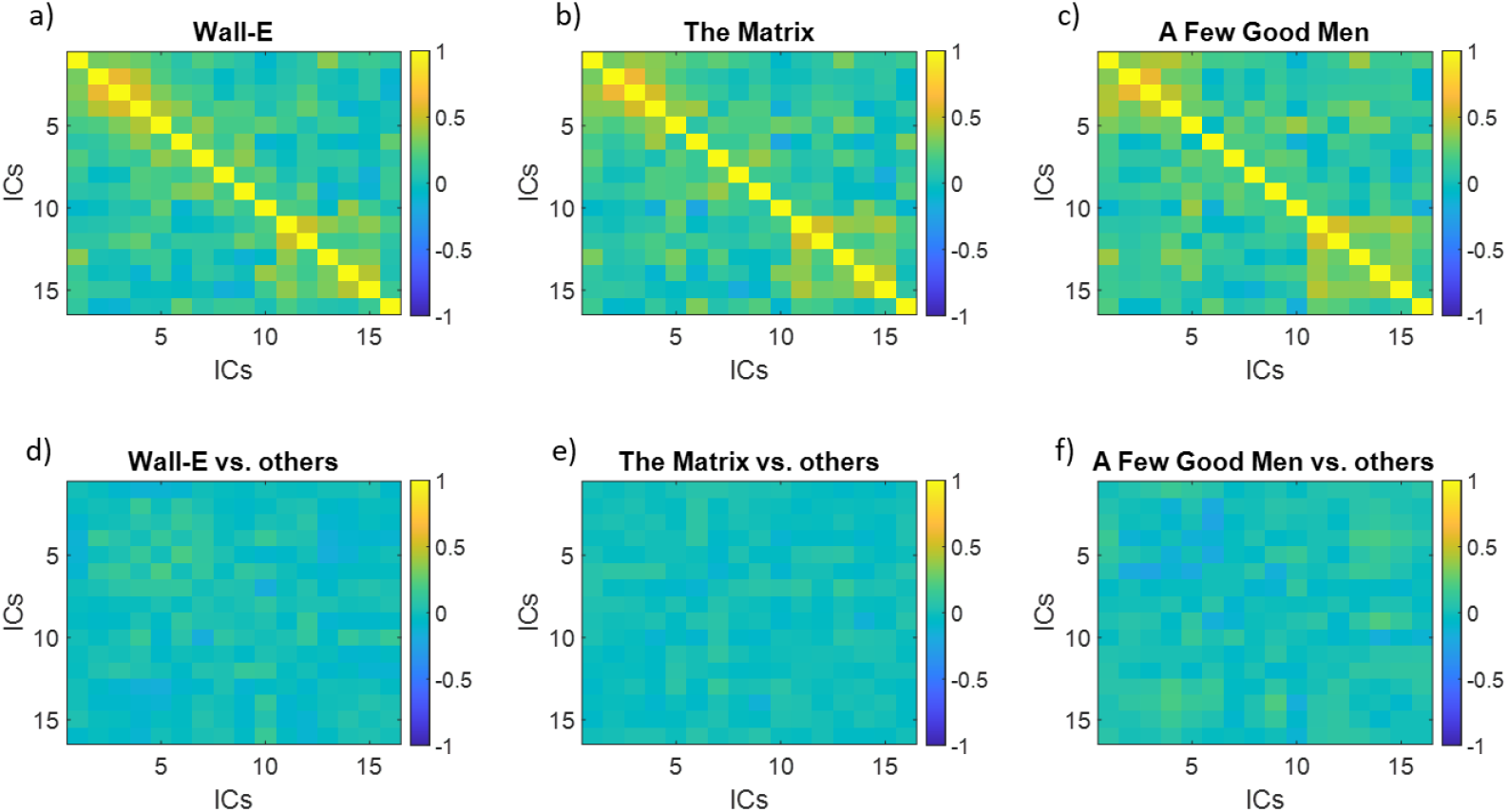
Top row, stationary connectivity among 16 networks (independent components, ICs) when watching the three movie clips from Healthy Brain Network Serial Scanning Initiative dataset. Bottom row, differences in stationary connectivity between a movie clip and the other two clips. No statistically significant difference was found even with an uncorrected threshold of p < 0.05.

### 3.2. Dynamic and stationary connectivity in 80-IC solution

We further studied the relations between dynamic and stationary connectivity in a larger spatial scale of the 80-IC solution. Figure 6 shows the dynamic connectivity consistency and stationary connectivity matrices for the four movie clips. The patterns are very similar to what was observed with the 20-IC solution. That is, the stationary connectivity matrices showed modular structures, and were very similar across different movie clips. In contrast, the dynamic connectivity distributions were highly skewed, with greater consistency between lower-level brain regions such as the visual and auditory cortex.

**Figure 6.**
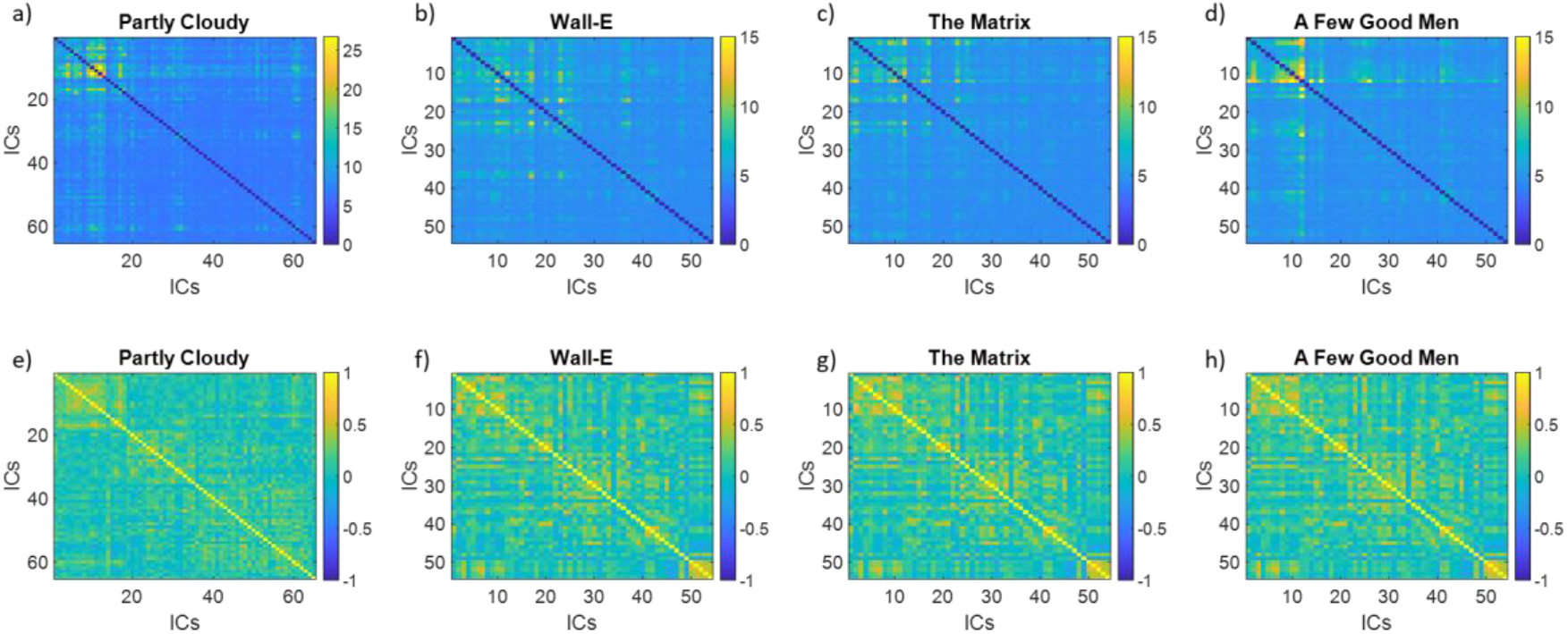
Dynamic connectivity consistency (top row) and mean stationary connectivity (bottom row) for the four movie clips using the 80-independent-component solutions. Please note that the number of included independent components (ICs) are different between the Partly Cloudy dataset and the other three movie clips.

We next directly examine the correlations among the matrices. For the Partly Cloudy data, the correlation between stationary and dynamic connectivity consistency was only 0.09 (Figure 7a), although it was statistically significant due to the large number of IC pairs. The relations have been confirmed by the HBN-SSI dataset (Figure 7b). The stationary connectivity of the three movie clips had almost perfect correlations (*ρs* = *0.97, 0.93,* and *0.95*). The dynamic connectivity of the three movies had weaker correlations (*ρs* = *0.70*, *0.62,* and *0.60*). In contrast, there were very small correlations between the dynamic and stationary connectivity matrices (*ρs* ranging from *0.23* to *0.28*).

**Figure 7.**
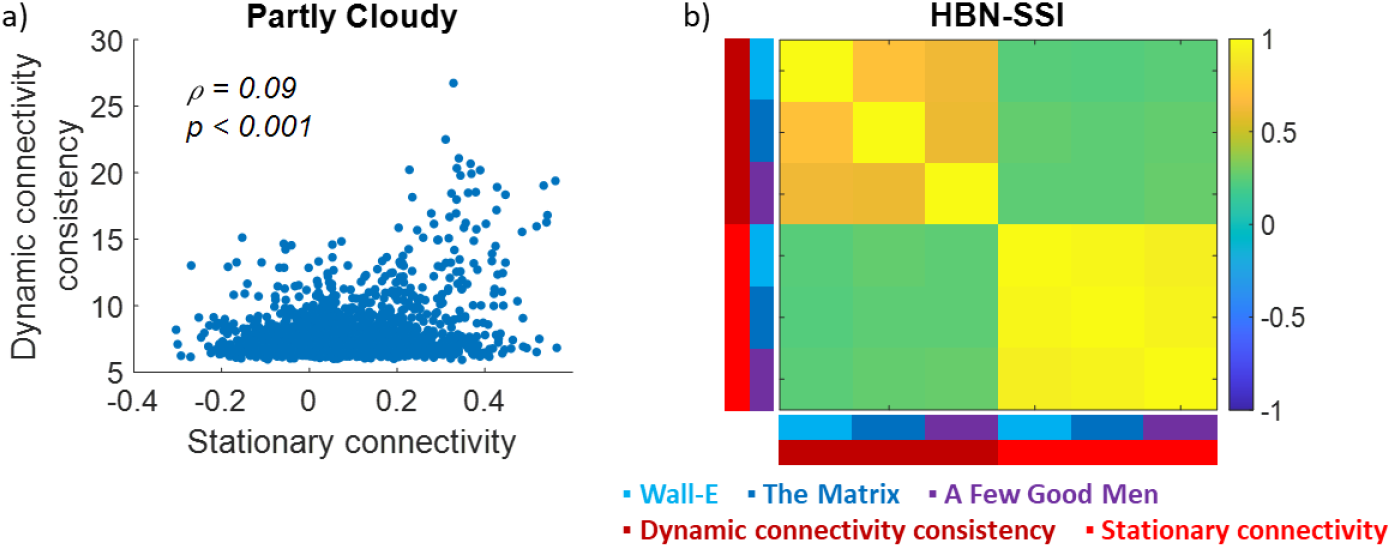
Correlations between stationary and dynamic connectivity for the Partly Cloudy dataset (a) and the Healthy Brain Network Serial Scanning Initiative (HBN-SSI) dataset (b). The connectivity matrices were calculated based on 80-independent-component solutions. Spearman’s rank correlation (ρ) was used.

To show the spatial distributions of dynamic connectivity in the context of global stationary connectivity, we calculated connectivity gradients based on the stationary connectivity of the three movie clips in the HBN-SSI dataset (Top row in Figure 8). The first and second gradients represented unimodal to transmodal gradient and visual to motor gradient, respectively. Next, we plotted the top 10% of dynamic connectivity in each movie clip compared with the other two movie clips (Bottom row in Figure 8). It can be seen that the consistent dynamic connectivity for the three movies usually took place between networks from far connectivity space, connecting visual, sensorimotor, and higher-order association systems. There are also notable differences among the three movie clips. For example, for the movie clips A Few Good Men, the consistent dynamic connectivity connected the higher-order associate areas to visual and sensorimotor regions, separately. But direct connections between visual and sensorimotor regions were rare.

**Figure 8.**
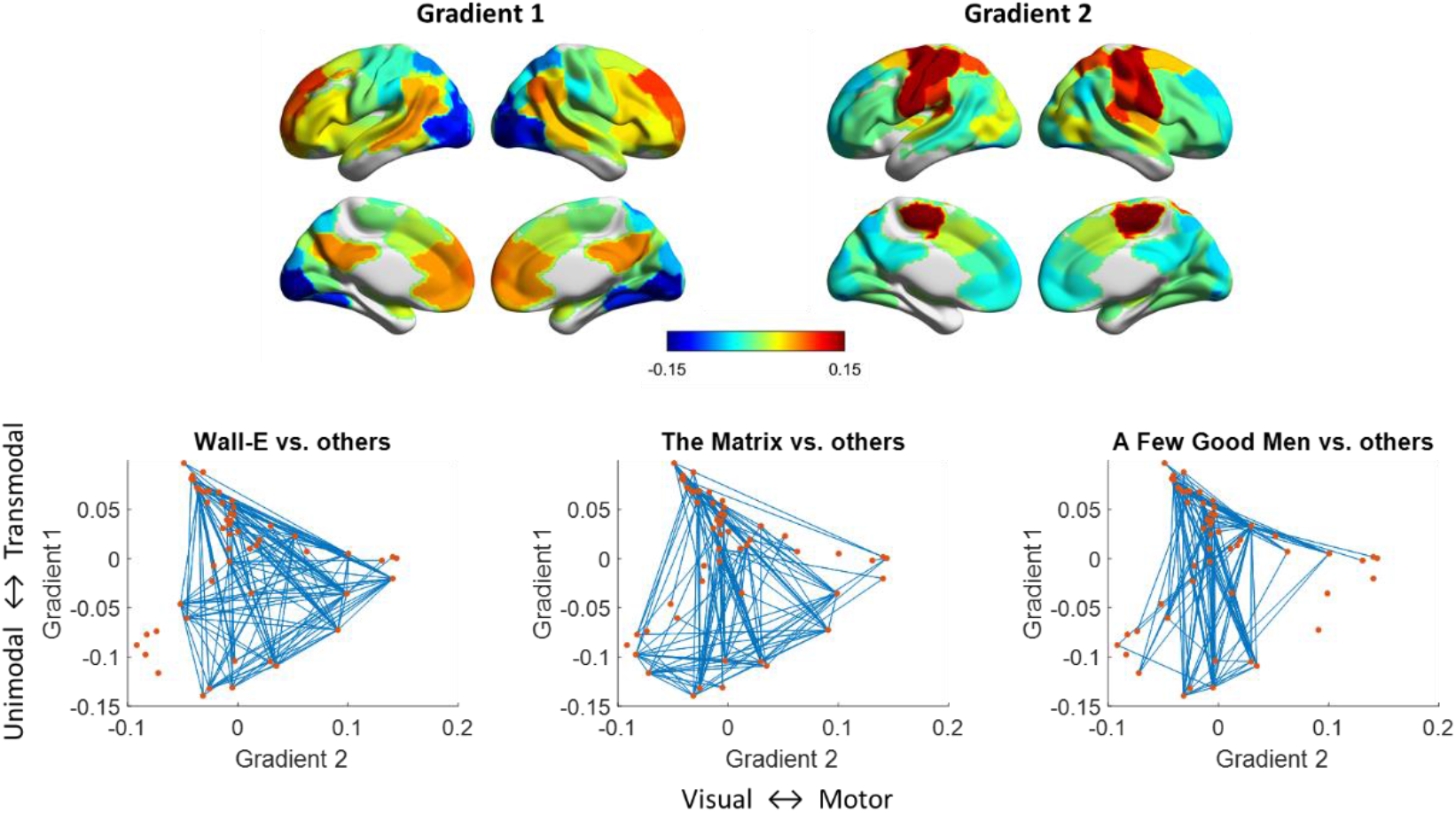
Top row, gradient maps of stationary connectivity in the Healthy Brain Network Serial Scanning Initiative dataset. Bottom row, top 10% consistent dynamic connectivity in each movie clip compared with the other two clips mapped to the connectivity gradient space.

### 3.4. Movie clips classification

Last we asked whether the dynamic or stationary connectivity pattern can enable individual-level prediction of the different movie clips. For each participant from the HBN-SSI dataset, we classified one of the three movie clips based on different measures. Overall, the consistency of dynamic connectivity achieved the highest prediction accuracy (*Accuracy* = 92.6%), followed by the stationary connectivity (*Accuracy* = 85.2%) and the consistency of regional activity (*Accuracy* = 74.1%). Compared with chance level accuracy of 33.33%, all classification accuracies were statistically significant (*p* < 0.001) based on permutation tests. The clip-to-clip classification results for the different features are shown in Supplementary Table S1.

## 4. Discussion

In this analysis, we have demonstrated widespread dynamic connectivity that was consistent across individuals when they watched the same movie clips. Different movie clips showed distinct patterns of dynamic connectivity, suggesting that the moment-to-moment interactions between brain regions may reflect the processing of context-specific information. For example, the two cartoon movie clips showed similarly consistent dynamic connectivity between the posterior cingulate and supramarginal networks. The clip of The Matrix showed more consistent dynamic connectivity in networks related to attention. And the drama clip, A Few Good Men, showed more consistent dynamic connectivity involving networks related to language processing, including bilateral fronto-parietal networks and prefrontal cortex. In contrast, the stationary connectivity showed very similar spatial patterns in different movie clips, with few statistical differences. The dynamic connectivity connected brain regions that are farther in the connectivity gradient space, and can better classify different movie clips than the stationary connectivity and regional activity.

We first empirically compared the statistical results of inter-individual correlations of dynamic connectivity measured by sliding-window and point-by-point multiplication. The multiplication approach showed higher statistical sensitivity than the sliding-window approach, as indicated by a much larger number of significant effects. It is not surprising because the point-by-point multiplication has kept the information of every time point, while the sliding-window approach can be seen as smoothed time series that could potentially filter out real signals. More specifically, the point-by-point multiplication approach detected consistent interactions between almost every pair of network ICs. This is in line with previous studies of regional activity, which also showed statistically significant effects in almost all cortical regions (Chen et al., 2016; Di and Biswal, 2020). In contrast, the sliding-window approach can only detect a small number of the dynamic connectivity among higher visual networks, and between supramarginal gyrus and posterior cingulate networks. This is probably because there is slow time-varying dynamic connectivity between these regions (e.g., Figure 1), which can be detected by the sliding-window approach. This is in line with studies showing that higher-order brain regions process longer time scale information (Baldassano et al., 2017). The results confirm the limitation of the sliding-window approach in studying dynamic connectivity.

Given the complexities of the video stimuli, it is difficult to pinpoint specific cognitive and affective functions involved in the different movie clips. But there are notable differences among them. The Wall-E clip involves only one character of a non-human robot, and the entire clip doesn’t have any conversations. In contrast, the other two clips involve human actors. And especially for the clip of A Few Good Men, there are intensive conversations among the characters. The clips may also differ in visual and auditory features, and other higher order features. Mapping these features is complicated, and may require additional techniques, such as computational models (Bartels et al., 2008; Nishimoto et al., 2011) or human ratings (Sun et al., 2022).

In the current results, we observed widespread consistent dynamic connectivity, but different movies clips were associated with distinct patterns of dynamic connectivity. The Wall-E clip showed a similar dynamic connectivity pattern as the other animated movie Partly Cloudy. Specifically, consistent dynamic connectivity was mainly observed in connectivity with the supramarginal and posterior cingulate networks. These regions involve higher-order social processes such as empathy and theory of mind (Richardson et al., 2018; Schurz et al., 2021). This makes sense because understanding the cartoon movies requires understanding the social interactions and intentions of the virtual characters. In contrast, the court drama clip, A Few Good Men, showed higher dynamic connectivity consistency that involved the auditory and temporoparietal junction networks. Because the clip mainly includes conversations among the characters, it is not surprising that the auditory cortex dynamically interacts with other cortical areas to pass auditory information to those areas. The temporoparietal junction is thought to be responsible for attributing other’s mental states (Koster-Hale and Saxe, 2013; Wang et al., 2021), which may also be a key component in understanding the conversations in the movie clip. Of course, these brain areas also involve in many other higher-order brain functions. For example, the supramarginal gyrus may also involve in spatial attention and working memory process (Silk et al., 2010). There might be alternative explanations on the cognitive functions involved in these networks during the watching of the clips. The correlational nature of the analysis doesn’t allow to reversely infer a specific function to the observed dynamic connectivity pattens (Poldrack, 2006).

In contrast, no differences were identified in stationary connectivity among the three movie clips. In a larger spatial scale of 54/80 ICs, we further showed that the spatial distribution of stationary connectivity was highly correlated among the three movie clips, which is consistent in previous studies (Tian et al., 2021; Vanderwal et al., 2019). The largely similar patterns of connectivity during watching different movies are also in line with observations in conventional task fMRI. When regressing out task activations (Cole et al., 2014; Di and Biswal, 2019) or using continuous task design (Krienen et al., 2014), the stationary connectivity or task-independent connectivity showed largely similar spatial patterns with each other and with what in resting-state. Similarly, the absolute correlation patterns of trial-by-trial variability in a stop signal task also showed similar patterns with each other and with a separate resting-state run (Di et al., 2020). Taken together, all the results convergently suggest that there is an overall connectivity pattern that may be related to the baseline brain function, but may also be related to the underlying physiology (Chen et al., 2020) or anatomical network structures (Laumann and Snyder, 2021). The lack of specificity of this global connectivity pattern makes it less desirable as a measure of brain connectivity in specific cognitive and mental conditions. It should be noted that some strategies may be used to extract stimuli related signals, e.g., retaining inter-individual consistent signals by using intersubject correlations (Di and Biswal, 2020; Simony et al., 2016). Simony and colleagues have proposed an intersubject functional connectivity approach to calculate stationary connectivity based on the shared signals across subjects. They have shown that the intersubject functional connectivity varied substantially between resting-state and story listening conditions (Simony et al., 2016). Based on similar strategy, it is possible to observe stronger differences in stationary connectivity among different movie clips than the conventional stationary connectivity methods used in the current analysis.

The spatial distributions of dynamic connectivity and stationary connectivity are largely dissociated. On one hand, the dynamic connectivity could take place between regions within the same functional systems, e.g., the visual system. It is reasonable that the lower-level sensory regions showed consistent interactions, but it can only be observed by using the point-by-point approach. This is less apparent when using the sliding-window approach, probably due to that the sliding-window approach can only capture slow fluctuations of dynamic connectivity. On the other hand, the current results showed that dynamic connectivity could also take place between different functional systems, e.g., between visual areas and the default mode network. This is in line with the economic account of brain network organizations, which suggests that transient communication between remote brain regions could enable efficient information transmissions. When overlaying the dynamic connectivity on the connectivity gradients space, it demonstrated more clearly that the dynamic connectivity took place between higher association areas, such as the default mode network, and lower-level sensory or motor regions. It should be noted that when using multivariate classification analysis, the spatial patterns of stationary connectivity can still be used to identify different movie clips, but with less accuracy than the dynamic connectivity patterns. The high classification accuracy of the dynamic connectivity suggested that dynamic connectivity could potentially be useful in predictive modeling analysis to reflect individual differences (Finn et al., 2015).

In the current study, we adopted a data-driven ICA approach to define regions of interest for the connectivity analysis. Interestingly, the components of interest, including the supramarginal and temporoparietal junction networks, covered left and right homotopic regions. The bilaterality still holds for the 80-IC solution. This suggests that the left and right homotopic regions in these networks are highly correlated. However, studies have suggested that the connectivity and functional activations of these regions may exhibit lateralization (Cai et al., 2013; Di et al., 2014; McAvoy et al., 2016). It is possible that the connectivity of these regions also showed lateralization effects, but were overlooked in the current analysis. Future studies may need to define unilateral regions of interest, if lateralization is an effect of interest.

One limitation of the current study is the sample size. The HBN serial scanning dataset has a relatively small sample size (n = 9). However, each participant watched three different types of movie clips and repeated four sessions, which enable us to directly compare connectivity among these diverse movie types and ensure the robustness of the results within an individual. Although promising, a larger sample size with an examination of behavioral scores is needed for future brain-behavioral association studies (Finn and Bandettini, 2021), where researchers can directly examine the differences in behavioral correlates between dynamic connectivity and stationary connectivity (Eichenbaum et al., 2021). Secondly, the current analysis only focused on the spatial distributions of dynamic connectivity. Given the ubiquitous dynamic connectivity identified in the current analysis, future studies could also examine the time courses of the point-by-point multiplications, which could paint a more complete picture of the dynamic connectivity.

## 5. Conclusion

By analyzing the inter-individual consistency of point-by-point multiplications between brain regions, we were able to identify functionally meaningful dynamic connectivity during movie watching. We found that compared with the stationary connectivity, the dynamic connectivity can be more sensitive to detect functional changes due to different movie contexts. The spatial distributions of dynamic connectivity and stationary connectivity were largely dissociated, with the dynamic connectivity reflect more long-range communications. Overall, the dynamic connectivity may provide more functionally relevant information than the stationary connectivity.

## Statements and Declarations

### Author Contributions

X.D., contributed to the study conception and data analysis. All authors have discussed the results. The first draft of the manuscript was written by X.D. and all authors commented on previous versions of the manuscript. All authors read and approved the final manuscript.

### Data availability

This study is a secondary analysis of publicly available datasets. The download links for the datasets are provided in the manuscript. No personal identifiable information was used in the current study.

