## Supplementary Materials for "Dynamic and stationary brain connectivity during movie watching as revealed by functional MRI"

S1. Intra-individual consistency is driven by the overall consistent components

S2. Differences in consistency of regional activity

S3. Sliding-window and point-by-point multiplication

S4. Individual-level movie clip classification

**S1. Intra-individual consistency is driven by the overall consistent components**

When different individuals watch the same movie stimuli, the brain responses in a certain brain region, *x*, can be partitioned into three components (Nastase et al., 2019), a common response across individuals, *c*, an idiosyncratic response for each individual, *id*, and the noise, *ε*.

$$x_{i}\left( t \right)=c\left( t \right)+{id}_{i}\left( t \right)+\varepsilon_{i}(t)$$

The common response, *c*, is what typically interested in, which can be measured by inter-individual correlation (Nastase et al., 2019) or principal component analysis (PCA) (Di and Biswal, 2021). The idiosyncratic response, *id*, represents the unique response pattern for each individual, which could be the similar when the same individual watch the same video multiple times.

For the Healthy Brain Network Serial Scanning Initiative dataset, each participant watched the same movie clips four times in separate scan sessions. Ideally, the multi-session and multi-participant design can be used to differentiate the consistent and idiosyncratic responses. We have explored whether there was significant idiosyncratic response on top of the common cross-participant consistent response. For the 16 independent components (ICs) from the 20-IC solution, we first calculated the inter-individual consistency of regional activity across all the 36 session/participants (Figure S1A). It can be seen that unimodal networks such as visual networks (IC # 1 through 4) and auditory network (IC # 6) had higher consistency compared with the other networks. Next, for each participant, we calculated the inter-session consistency of regional activity across the four sessions. Although noisier, we could still find similar network ICs had that higher inter-session consistency across participants (Figure S1B). We then regressed out the overall consistent response calculate from the entire 36 session/participants, and then calculated the inter-session consistency for each participant based on the residual time series (Figure S1C). It can be seen that all the network ICs and participants seemed to have similar level of consistencies, suggesting that they were at chance level. Therefore, we conclude that individually consistent idiosyncratic responses across sessions were not apparent in the 16 networks’ regional activity time series. Moreover, the idiosyncratic responses are not the focus of the current study, because the focus is to examine the differences of common responses among different movie clips. Therefore, in the current analysis, we treated session and participant equally as separate data and calculated inter-individual consistency across all the sessions and participants.

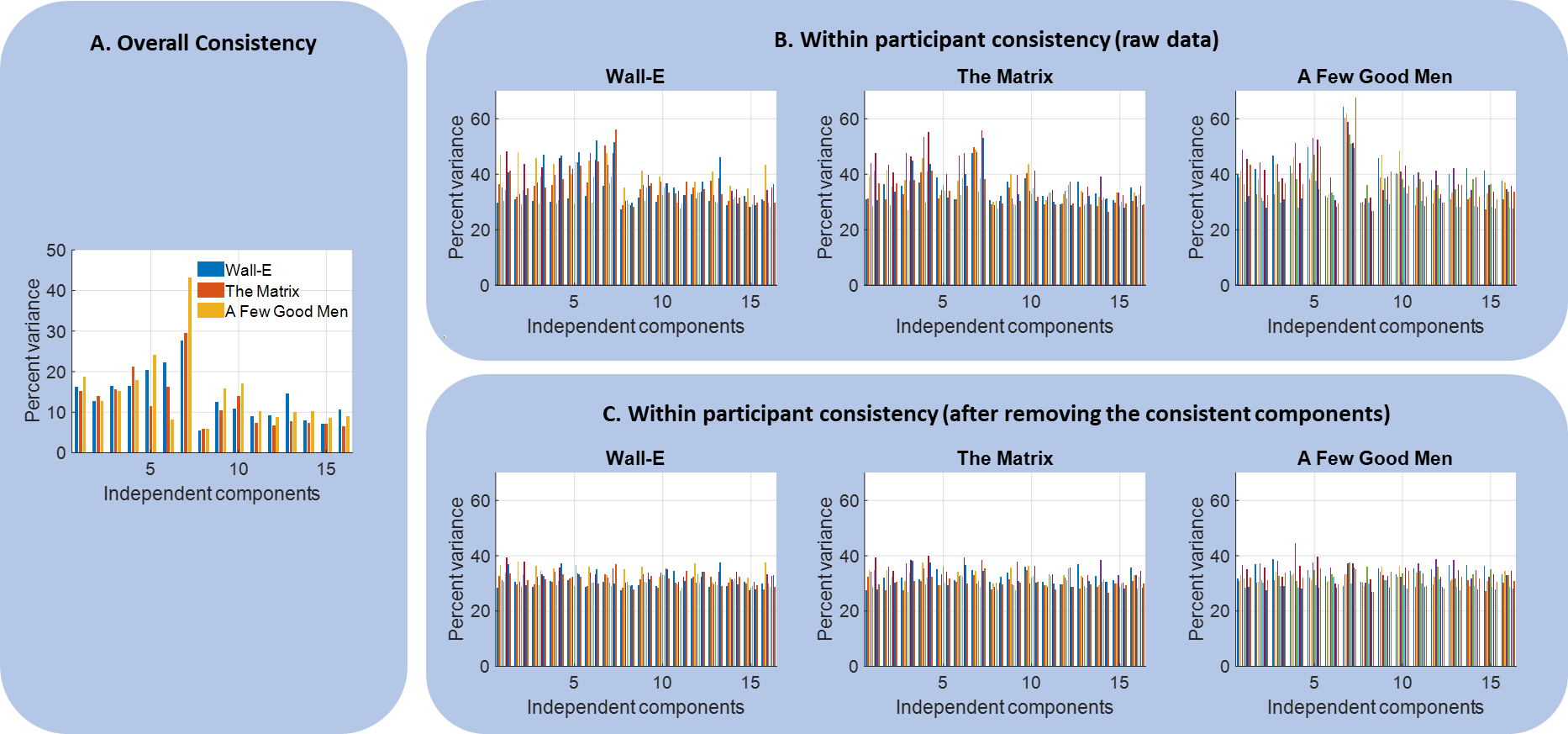

**Figure S1**

**S2. Differences in consistency of regional activity**

For the HBN-SSI, we further examined whether the variances explained by the first PCs are different among the three movie chips. We compared the variance explained by the first PC in one movie clip with the averaged variance explained by the first PCs of the other two clips. We performed permutation tests to determine the statistical significance. Specifically, we had three 410 x 36 matrices *X* for the three movies. Permutation was performed by randomly assign the 108 (36 x 3) time series into three new 410 x 36 matrices. PCA was performed separately for the three randomized matrices, and the differences between the variance in one matrix between the averaged variance of the other two matrices was obtained. The permutation was performed 10,000 times, resulting in 30,000 randomized values of differences in variance between one matrix and the other two matrices. The differences in variance explained by each movie compared with the other two movies were compared with the null distribution of obtain p values. FDR correction of p < 0.05 was used.

The networks (ICs) that showed different consistency of regional activity among the three movie clips are shown in Figure S2. Four networks (ICs) showed higher inter-individual correlations in Wall-E compared with the other two movie clips, including the posterior cingulate cortex, supramarginal gyrus, left fronto-parietal, and medial and lateral prefrontal networks. Only one network covering the posterior parietal lobe showed higher inter-individual synchronization in The Matrix compared with the other two movie clips. Lastly, eight networks showed higher inter-individual correlations in A few Good Men compared with the other movie clips, including the auditory cortex, medial visual, temporo-parietal junction, and a few fronto-parietal networks.

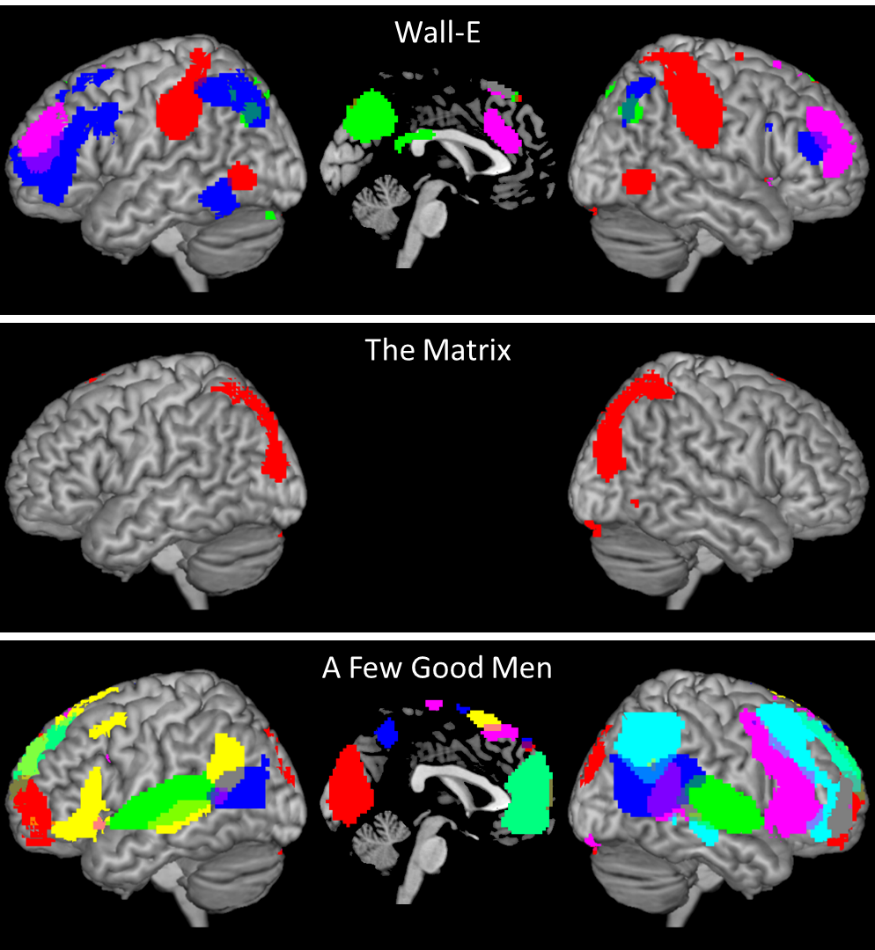

**Figure S2** The brain networks (independent components, ICs) that showed different synchronization between the three movies from the healthy brain network serial scanning initiative. The IC maps represent higher inter-individual consistency in one movie clip compared the other two clips (p < 0.05 false discovery rate with permutation test).

**S3. Sliding-window and point-by-point multiplication**

The sliding-window approach is the most commonly used method to estimate dynamic connectivity. We performed sliding-window analysis using a similar setting as our previous work (Di and Biswal, 2020). Specifically, we used a 30 TR (60 s) window with a window step of 1 TR (2 s). For each pair of networks (ICs) in a participant, Pearson’s correlation coefficient was calculated for each window and transformed into Fisher’s z scores, forming a dynamic connectivity time series. PCA was performed on the time-window by participant matrix (139 x 29) to obtain variance explained by the first PC. To determine the statistical significance of the variance explained by the first PC, we performed circular time-shift randomization to determine the null distribution (Di and Biswal, 2021; Kauppi et al., 2010). The time series from the two network ICs from all the participants were circular shifted with random delays. Sliding-window dynamic connectivity were then calculated for each participant, and PCA was performed. The randomization was performed 10,000 times for each pair of networks. The real values were compared with the null distribution to perform statistical inferences. This resulted in a 16 x 16 matrix. FDR correction was used to correct for multiple comparisons (120: 16 x 15 / 2). Because theoretically the sliding-window should have less signal to noise ratio than the point-by-point multiplication approach. And the analysis on the Partly Cloudy dataset has confirmed this. The sliding-window approach was not performed for the remaining analyses.

We first compared the two methods for dynamic connectivity estimation on the Partly Cloudy dataset. Figure S3 shows their inter-individual consistency among the 16 networks (ICs) from 20-IC solution. Four pairs of networks (ICs) showed consistent dynamic connectivity using the sliding-window method (Figure S3a), mainly among higher visual regions extending to the posterior parietal network and supramarginal network (IC # 4 and 6). One additional connectivity was between the supramarginal and default mode network (IC # 6 and 15), which was similar to our previous analysis using seed-based approach. In contrast, the point-by-point multiplications were more widespread, with the highest consistency among the higher visual networks. This makes sense because the point-by-point multiplication retains more information about the dynamics compared with the sliding-window approach. Therefore, in the following analysis, we only used the multiplication approach to study dynamic connectivity.

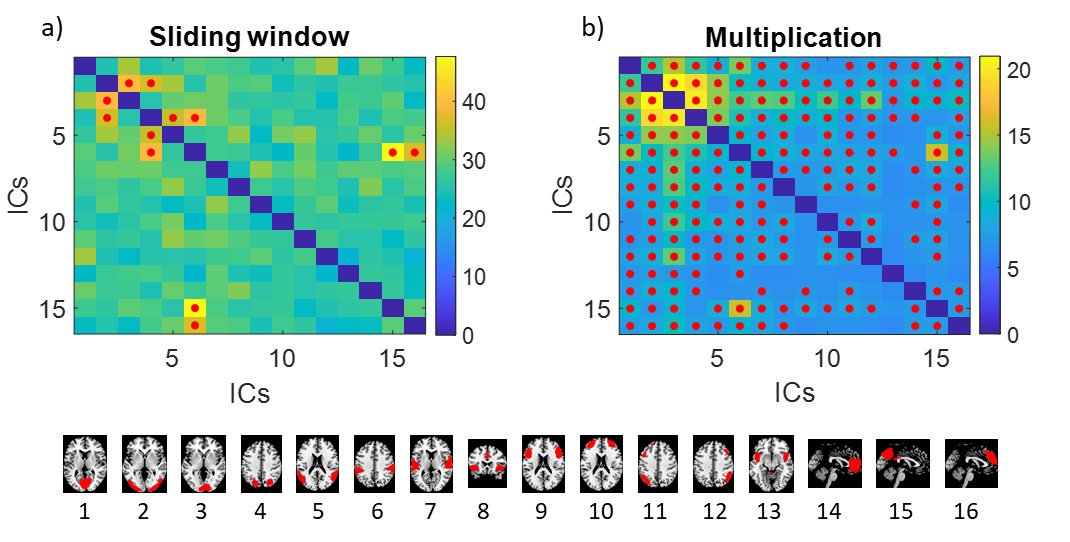

**Figure S3** Inter-individual consistency of sliding-window dynamic connectivity (a) and point-by-point multiplication (b) from the Partly Cloudy dataset. The matrices were calculated across 16 networks (independent components, ICs), which are shown at the bottom. The red dots indicate false discovery rate (FDR) p < 0.05 using a circular time-shift randomization procedure.

**S4. Individual-level movie clip classification**

|  |  | Predicted | | |
| --- | --- | --- | --- | --- |
|  |  | Wall-E | The Matrix | A Few Good Men |
| Dynamic connectivity | | |  |  |
| Input | Wall-E | 0.78 | 0.11 | 0.11 |
|  | The Matrix | 0.00 | 1.00 | 0.00 |
|  | A Few Good Men | 0.00 | 0.00 | 1.00 |
| Stationary connectivity | |  |  |  |
| Input | Wall-E | 0.78 | 0.22 | 0.00 |
|  | The Matrix | 0.00 | 0.89 | 0.11 |
|  | A Few Good Men | 0.11 | 0.00 | 0.89 |
| Regional activity | |  |  |  |
| Input | Wall-E | 0.67 | 0.22 | 0.11 |
|  | The Matrix | 0.11 | 0.78 | 0.11 |
|  | A Few Good Men | 0.00 | 0.22 | 0.78 |

**Table S1** Confusion matrices for the movie classifications based on dynamic connectivity, stationary connectivity, and regional activity patterns.
